# Generic solving of physiologically-based kinetic models in support of next generation risk assessment due to chemicals

**DOI:** 10.1101/2022.04.29.490045

**Authors:** Sandrine Charles, Ophelia Gestin, Jérémie Bruset, Dominique Lamonica, Virgile Baudrot, Arnaud Chaumot, Olivier Geffard, Thomas Lacoue-Labarthe, Christelle Lopes

**Affiliations:** University of Lyon, University Lyon 1, UMR CNRS 5558, Villeurbanne, France; INRAE, Riverly, Ecotoxicology, Lyon, France; University of La Rochelle, UMRi 7266, La Rochelle, France; Qonfluens, Rdpt Benjamin Franklin, Montpellier, France

**Keywords:** Pharmacokinetics, Toxicokinetics, General unified modelling framework, Bioaccumulation, Chemical exposure

## Abstract

Increasing the confidence in using *in vitro* and *in silico* model-based data to aid the chemical risk assessment process is one, if not the most, important challenge currently facing regulatory authorities. A particularly crucial concern is to fully take advantage of scientifically valid Physiologically-Based Kinetic (PBK) models. Nevertheless, risk assessors remain still unwilling in employing PBK models within their daily work. Indeed, PBK models are not often included in current official guidance documents. In addition, most users have limited experience in using modelling in general. So, the complexity of PBK models, together with a lack to evaluation methods of their performances, certainly contributes to their under-use in practical risk assessment.

This paper proposes an innovative and unified modelling framework, in both the writing of PBK equations as matrix ordinary differential equations (ODE), and in its exact solving simply expressed with matrix products. This generic PBK solution allows to consider as many as state-variables as needed to quantify chemical absorption, distribution, metabolism and excretion processes within living organisms when exposed to chemical substances. This generic PBK model makes possible any compartmentalisation to be considered, as well as all appropriate inter-connections between compartments and with the external medium.

We first introduce our PBK modelling framework, with all intermediate steps from the matrix ODE to the exact solution. Then we apply this framework to bioaccumulation testing, before illustrating its concrete use through complementary case studies in terms of species, compounds and model complexity.

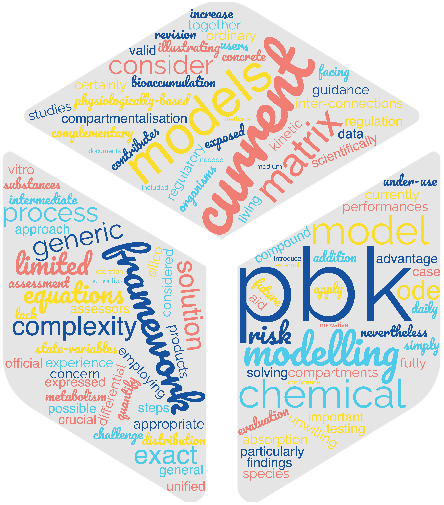

## 1 Introduction

Physiologically-Based Kinetic (PBK) models encompass both Physiologically-Based Pharmaco-Kinetic (PBPK) and Physiologically-Based (eco-)Toxico-Kinetic (PBTK) models depending on the context referring either to therapeutic drug development (Zhang et al., 2006) or environmental risk assessment (Grech et al., 2017; Mit et al., 2022). All PBK models are compartment models employing ordinary differential equations (ODE) to quantify chemical absorption, distribution, metabolism (i.e., biotransformation) and excretion (ADME) processes within living organisms when exposed to chemical substances (Wang and Rainbow, 2008). One fundamental aspect of PBK model complexity is the degree of compartmentalization (i.e., differentiation of an organism into various tissues or organs) (Armitage et al., 2021). This complexity translates into a high number of parameters that are usually valued from literature information or expert knowledge. As a consequence, most of PBK models have purely predictive usages, for example for human health assessments where novel experiments are very limited, even impossible (OECD, 2021). Nevertheless, recent advances in the development of Bayesian inference tools make today possible to deal with complex models (Fransson et al., 2014; Sweeney et al., 2015; Testai et al., 2021; Charles et al., 2021) even with sparse data sets, which opens up new possibilities for PBK models to obtain parameter estimates associated with their uncertainties and to propagate this information to model-informed predictions. Also, as part of New Approach Methodologies (NAM), PBK models as well as *in vitro-in vivo* extrapolation (IVIVE) approaches can be combined with bioactivity data, all of this being helpful for prioritizing thousands of “data poor” chemicals in human health risk assessment (Armitage et al., 2021; Breen et al., 2021).

Relying *a priori* on the anatomical and physiological structure of the body, PBK model compartments usually correspond to target organs or tissues, possibly all interconnected and related to the external exposure medium. However, lots of PBK models only relate organs or tissues to blood or lymph, with only one or two organs or tissues linked to the external medium (Zhang et al., 2019, 2022). Unfortunately, such simplifications of PBK models are often justified based on technical bases rather than physiological ones. All PBK models are written as coupled ODE whose parameters correspond to fluxes between compartments, for which information is partly available in the scientific literature. The general tendency to develop more complex models to include as many physiological processes as possible, in addition to the current computational capacities, may give the impression that the most complex models will be the most efficient. Nevertheless, with an increasing number of parameters to calibrate, such complex models may rather be avoided in favour of a balance between complexity and simplicity. All the more so since, as a matter of fact, current regulatory documents for assessing the bioaccumulation of chemicals in organisms usually only require simple one-compartment models (Organisation for Economic Co-operation and Development, 2012; Ratier et al., 2019, 2021; Charles et al., 2021), even if it is now recognized that it is necessary to also consider internal concentrations within target organs in order to fully capture the chemical bioaccumulation behaviour, the specific role of organs, and the dynamics of toxic effects (Vijver et al., 2004).

Multi-compartment models reveal sometimes absolutely necessary, for example in order to finely decipher the internal contamination routes of specific chemical compounds causing damages to only specific organs (Brinkmann et al., 2014; Allen and Weihrauch, 2021; Gestin et al., 2021). Additionally, PBK models can be of crucial importance to predict organ-level concentration–time profiles in situation where animal testing is now prohibited, using PBK model information from one chemical substance to inform the development or evaluation of a PBK model for a similar chemical substance (Thompson et al., 2021). In the perspective to enlarge and facilitate the use of PBK models, namely to include more compartments, to better estimate parameter values from data and to better support a fine deciphering of underlying contamination processes after chemical exposure, there is today a clear need for user-friendly tools. From an automatized implementation, fully transparent and reproducible, such tools should simplify the use of any PBK models, preventing users to invest into technicalities, whatever the required number of compartments to physiologically consider, whatever the number of connections to account for between compartments in pairs or between compartments and the external medium, and whatever the species-compound combination of interest. Such tools seem the only way to gradually achieve greater acceptability of complex PBK models, even in a regulatory context (Paini et al., 2019; Armitage et al., 2021).

Capitalizing on recent publications on TK models (Ratier et al., 2021; Charles et al., 2021), we present in this paper a very innovative exact solution of a fully generic PBK model written as a set of ODE. Benefiting from an exact solution is a tremendous advantage in terms of numerical implementation. Indeed, it avoids to discretise ODE as there is no more numerical approximation, the solution being directly used for the inference process and the subsequent simulations. The gain in calculation time is huge (more than 100-fold), associated with a fair use of computer resources. Moreover, the new modelling framework we propose makes possible to account for an infinite number of compartments, with all possible connections between pairs of compartments and between compartments and the exposure medium, independently of the investigated species or chemical substance. Indeed, we found a particularly condensed way to write a linear ODE system, which is typical of PBK models, thus allowing us to fully and exactly solve the ODE system to write a generic exact solution. This exact solution reveals particularly interesting when estimating a large number of parameters related to many state variables and for which experimental data may be sparse, with few replicates and a high variability.

In the following sections, we first detail our generic modelling framework, together with notations of parameters and variables, providing the generic solution at the end of section 2. In section 3, the generic modelling framework is applied to the particular context of bioaccumulation of chemical substances within organisms. We detail how to write the ODE system for both accumulation and depuration phases of standard bioaccumulation tests, and then how to get the final generic solution to simulate internal concentrations over time from parameter estimates. Lastly, section 4 presents three different situations where simulations revealed useful to predict what happens within organs and/or tissues according to the species under consideration, and to the chemical substance to which it is exposed. These case-studies were chosen from the literature to be both diverse and complementary in terms of questions that a PBK model can help to investigate. We thus present to two case studies for the species *Gammarus fossarum* exposed to cadmium, for which internal concentrations have been measured within four organs: one case study with one-compartment PBK models for each organ considered separately; the other case study with a four-compartment PBK model. From these case studies we illustrate the added-value of considering one single four-compartment model rather than four one-compartment models for each organ. The third case study concerns the species *Danio rerio* exposed to arsenic, whose bioaccumulation process is described with a six organ-based compartment PBK model.

## 2 Generic modelling framework

Biologists will often have the tendency to expect an exhaustive description of phenomenon they are studying. In the same way, mathematicians will want to use the most sophisticated methods they know. However, all models are inherently wrong, only some of them will prove useful (Box, 1976). As a consequence, the modeller should position between these two points of view to be efficient. Such a position is known as the *parsimony principle* by which the simplest model that adequately explains the data should be used; it was proposed in the 14^th^ century by William of Ockam, an English Franciscan friar, scholastic philosopher, and theologian (Green et al., 2018). In its general form, the parsimony principle also referred to as Occam’s Razor; it states that the simplest of competing explanations is the most likely to be correct. In the context of model fitting, the simplest model providing a good fit will be preferred over a more complex one. The compromise is thus between good description of the observed data and simplicity.

In this spirit, the most generic PBK models will rely on simplifying hypotheses, to both decipher enough internal mechanisms under chemical exposure, and remain mathematically reasonable so as to be easily manipulated. Below is a non-prioritized short list of the most current hypotheses: (i) the exposure concentration is assumed constant over time; (ii) there can be any number of compartments in direct relation with any number of tissues and organs that are need to consider on a biological point of view; (iii) all compartments can be connected two-by-two to all the others; (iv) the exposure contaminant can enter within each compartment; and additionally, (v) all compartments can be theoretically connected to the external medium, the final choice to be based on biological expertise.

From these hypotheses, Figure 1 gives the general schematic representation of exchanges between the external medium and compartments, as well as between compartments themselves. Table 1 gathers all variables and parameters involved in the generic writing of the PBK model, and used in Figure 1.

**Figure 1:**
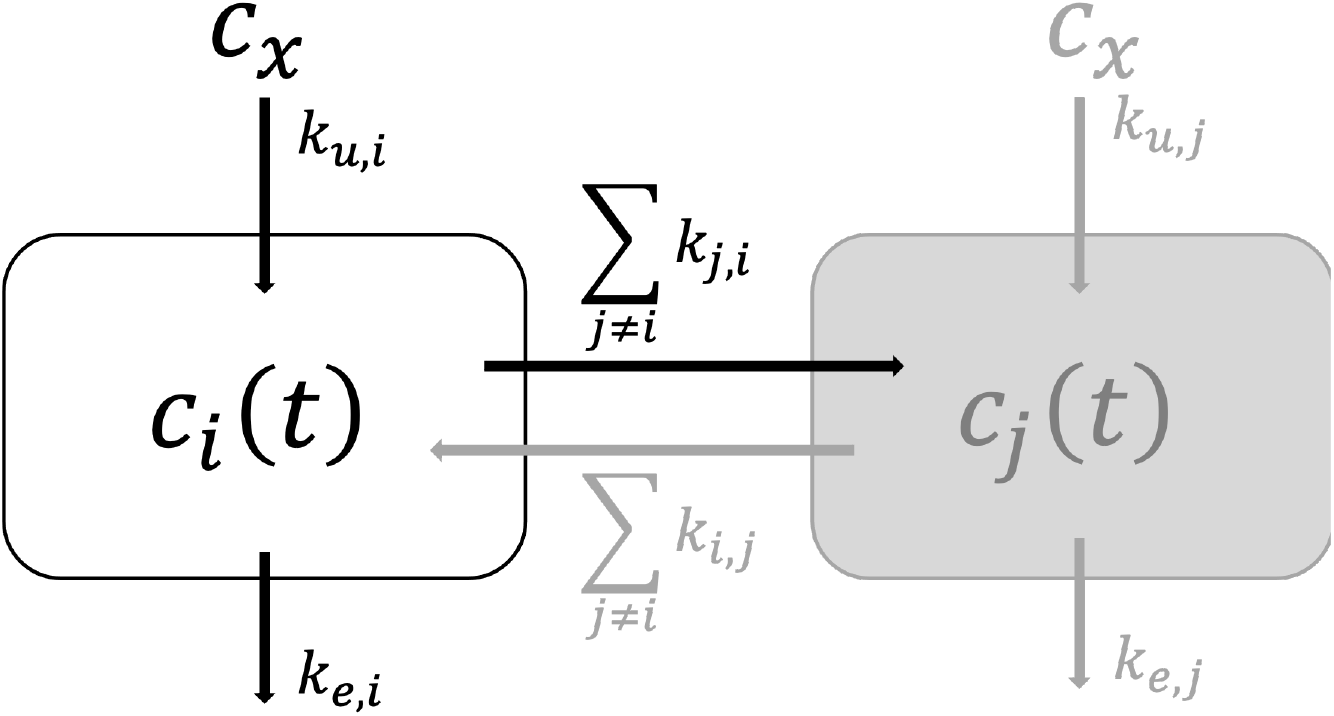
Generic scheme of a multi-compartments toxicokinetic model connecting *n* compartments two-by-two and each of them to the external medium at exposure concentration *c_x_* (*i, j* ∈ [1; *n*]). Refer to Table 1 for names, meaning and units of variables and parameters.

**Table 1:** Variable and parameter names, meaning and unit as used within the generic PBK model all a long this paper. Symbol # means dimensionless; [*t*] stands for time unit.

| Names | Meaning | Unit |
| --- | --- | --- |
| $t$ | time | $[t]$ |
| $n$ | total number of compartments | $\#$ |
| $i, j$ | Compartment numbers | $i, j \in [1; n]$ |
| $c_x$ | exposure concentration in the external medium | mass per volume |
| $c_i(t)$ | internal concentration in compartment $i$ at time $t$ | mass per weight |
| $k_{u,i}$ | uptake rate from the external medium to compartment $i$ | $[t]^{-1}$ |
| $k_{e,i}$ | elimination rate from compartment $i$ to the external medium | $[t]^{-1}$ |
| $k_{i,j}$ | input rate from compartment $j$ to compartment $i$ | $[t]^{-1}$ |
| $k_{j,i}$ | output rate from compartment $i$ to compartment $j$ | $[t]^{-1}$ |

### 2.1 Mathematical equations of the PBK model

From Figure 1, we can derive the full system of ordinary differential equation (ODE) describing the dynamics of the multi-compartments model when organisms are exposed to an external constant concentration *c_x_*:

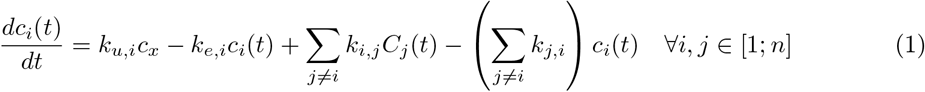

Names, meaning and units of variables and parameters are provided in Table 1.

This full system of ODE for *n* compartments all related by pairs (equation 1) can equivalently be written in a matrix way as follows:

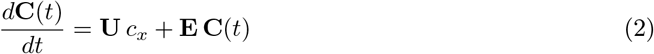

where vector **C**(*t*) gathers all internal concentrations in compartments *i* at time *t*, *i* ∈ [1; *n*]:

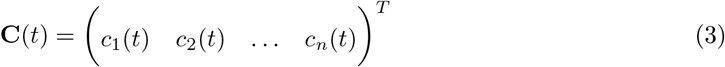

Vector **U** contains all uptake rates from the external medium at exposure concentration *c_x_*:

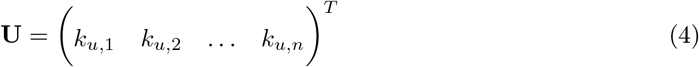

Matrix **E** gathers both input and output rates between compartments two-by-two, together with the elimination rates from each compartment *i, i* ∈ [1; *n*]:

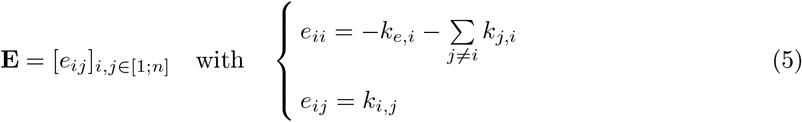

Equation (2) is a matrix ODE system, which is linear with a second member. It can be solved in two steps as detailed below.

### 2.2 Generic solving the PBK model

The first step is to solve the matrix system (2) without its second member. Then, the second step consists in finding the final general solution using the method of the variation of constant.

#### 2.2.1 Solving the ODE system without the second member

Removing the second member from the matrix ODE system (2) leads to the following system to solve:

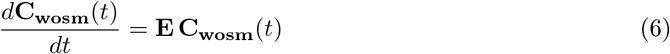

with **C_wosm_**(*t*) the desired solution of equation (6) without second member (abbreviated by index *wosm*). Using matrix exponential immediately provides the solution:

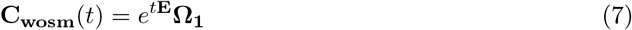

with **Ω**_1_ a vector integration constant 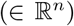, and the following definition for the matrix exponential:

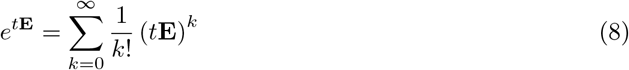

#### 2.2.2 Getting the generic solution of the ODE system

For the second step including the second member, we used the method of the variation of the constant, starting from the assumption that the general final solution with second member (abbreviated by index *wsm*) can be written as follows:

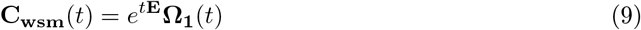

with a vector function **Ω**_1_ (*t*) to be determined.

Given that the exposure concentration is assumed constant (equal to *c_x_*) in this paper, deriving equation (9) and replacing terms in equation (2) leads to the following result:

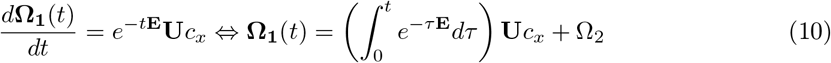

The final generic solution of equation (2) will thus write as:

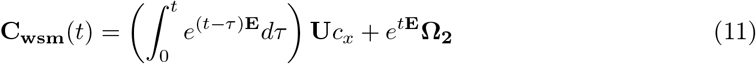

with 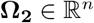 a constant to be determined.

From an initial condition **C_wsm_**(*t* = 0) = **C**_0_, we finally get **Ω**_2_ = **C**_0_, which leads to the following final particular solution of equation (2):

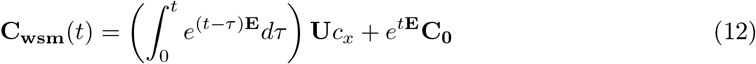

It remains now to calculate the matrix integral to achieve the very final solution of the matrix ODE system (2).

#### 2.2.3 Final expression solution of the PBK solution

As detailed in the Supplementary Information (SI), and using the definition of a matrix exponential from equation (8), the matrix integral in equation (12) can be calculated with the following expression:

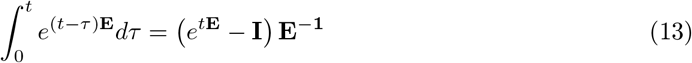

where matrix **I** is the identity matrix, *i.e.* the (*n* × *n*) square matrix with ones on the main diagonal and zeros elsewhere, and **E**^-1^ the inverse matrix of **E**.

It can immediately be deduced that the final solution in equation (12) simplifies as follows:

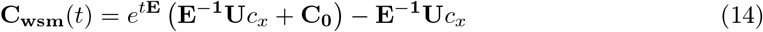

In the next sections, we employed this generic expression to go beyond and build generic physiologically-based kinetic models, that we applied to different case-study in the field of ecotoxicology.

## 3 Application to bioaccumulation testing

### 3.1 Accumulation and depuration phases

Bioaccumulation is defined as an increase in contaminant concentrations inside living organisms following uptake from the surrounding medium (living media, food, even workplace for humans). Bioaccumulation is the result of dynamical processes of uptake and elimination that can be modelled with the above ODE system (equation (2)). The extent to which bioaccumulation occurs within a given species determines the subsequent toxic effects. Hence, a better knowledge on bioaccumulation enables to assess the risk related with the exposure to chemicals and to evaluate our ability to control their use and emissions in the field (Chojnacka and Mikulewicz, 2014). Bioaccumulation is thus the net result of all uptake and elimination processes, by egestion, passive diffusion, metabolization, excretion and maternal transfer. Concomitantly, the organism growth modulates the bioaccumulation by diluting chemical quantities in increasing body or organ mass (Figure 2).

**Figure 2:**
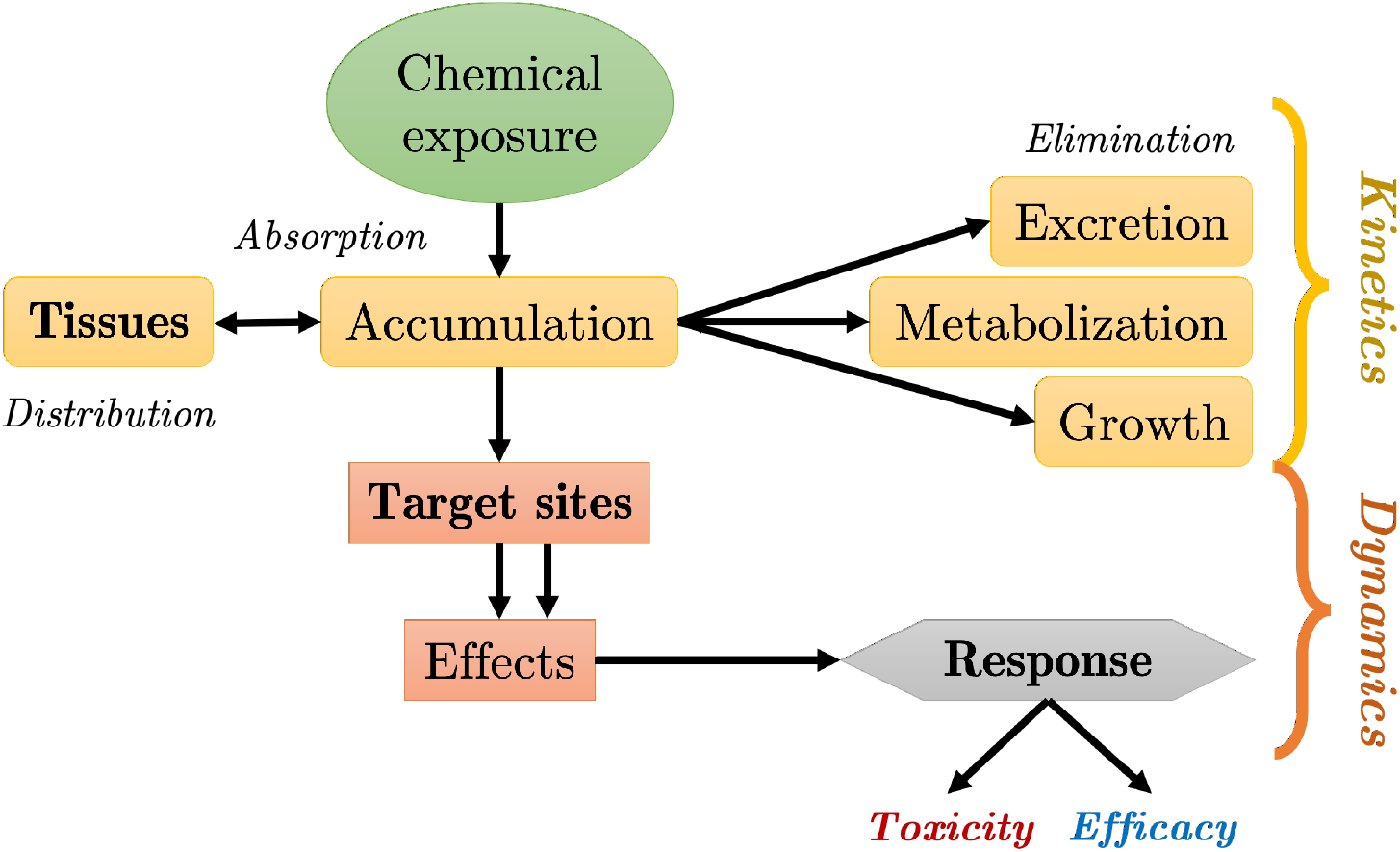
Absorption, Distribution, Metabolism and Excretion (ADME) processes and their relationships with effects and responses within living organisms, leading to either toxicity or efficacy depending on the type chemical substance to which they are exposed.

Bioaccumulation tests are usually mid-to-long term laboratory experiments designed to identify all the potential uptake pathways, including food and waterborne exposure routes (Marigomez, 2014). Bioaccumulation tests commonly comprise an accumulation phase followed by a depuration phase (European Commission, 2013). During the accumulation phase, organisms are exposed to a chemical substance of interest. After a certain time period (for *t* ∈ [0; *t_c_*]), with *t_c_* fixed by the experimental design, organisms are transferred to a clean medium for the depuration phase (for *t* > *t_c_*). The concentration of the chemical substance (and of its potential metabolites) within organisms is then measured internally at regular time points during both phases. In an ERA perspective, such data can be used *in fine* to estimate bioaccumulation metrics (Ratier et al., 2021).

If the bioaccumulation within organisms is widely studied for humans and large animals, namely, fish, birds, and farm animals (Tarazona et al., 2015; Grech et al., 2019), this is less the case for invertebrates (Amyot et al., 1996). However, it is equally important to decipher internal processes at the target organ level in invertebrates. Indeed, deeply unravelling internal routes of chemical substances between organs after they entered within the body, is a of great interest to better understand mechanisms implied in the subsequent effects on fitness, a phenomenon known as organotropism (Ju et al., 2011; Rocha et al., 2015; Chen et al., 2018). Among invertebrate species of interest, crustacean amphipods are already recognized particularly relevant as aquatic biomonitors of trace metals (Amyot et al., 1996; Gust and Fleeger, 2005; Ingersoll et al., 2005; Van Geest et al., 2010; Kuehr et al., 2021).

### 3.2 Generic modelling of bioaccumulation

Within this context, the generic ODE system (equation 2) may fully be applied to describe, simulate and predict what happens within organs (in terms of internal concentration over time) and between organs (in terms of uptake, elimination and exchange rates) when an organism is exposed to a given chemical substance. To this end, each organ can be associated with one model compartment, leading to the following equations for both accumulation and depuration phases:

**Accumulation phase** (0 ≥ *t* ≥ *t_c_*)

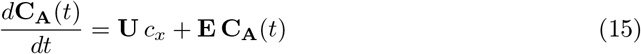 Equation (15) is identical to equation (2), denoting **C_A_**(*t*) the internal concentration at time *t* during the accumulation phase.

**Depuration phase** (*t* > *t_c_*)

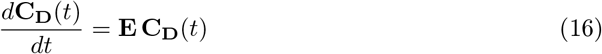

Variable **C_D_**(*t*) is the internal concentration at time *t* during the depuration phase. Parameters and variables have the same meaning as given in Table 1.

Regarding the accumulation phase, equation (15) has a solution directly given by equation (14), whatever the initial condition equal to **C_A_**(*t* = 0) = *C*_0_.

As a consequence, the generic solution for the **accumulation phase** writes as follows:

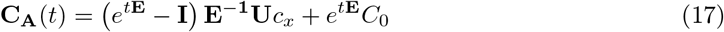

Regarding the depuration phase, equation (16) is similar to equation (6), with the corresponding generic solution given by equation (7). The constant vector **Ω**_1_ can be determined from the initial condition of the depuration phase that corresponds to the internal concentration reached at *t* = *t_c_* at the end of the accumulation phase. We must therefore solve the following equation:

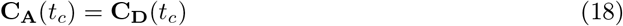

Given solutions from equations (14) and (7), we get:

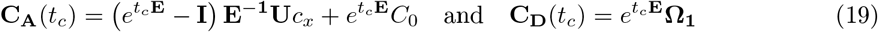

Hence, the constant vector **Ω**_1_ derives from the following equation:

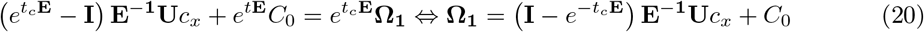

The generic solution for the **depuration phase** then writes as follows:

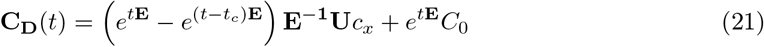

## 4 Case studies in ecotoxicology

This section presents three case studies with different numbers of compartments. Two case studies concern species *Gammarus fossarum* exposed to cadmium (Cd) as studied in Gestin et al. (2021), the third one concerns species *Danio rerio* exposed to arsenic (As). Gestin et al. (2021) used both several one-compartment TK models compared with one four-compartment TK model, to gain knowledge on the accumulation and fate dynamic of Cd in and between gammarids’ organs. Subsections 4.1 and 4.2 give the generic solutions of parameter estimates.

### 4.1 Case study with one compartment

Applying equation (2) with only one compartment leads to two single equations:

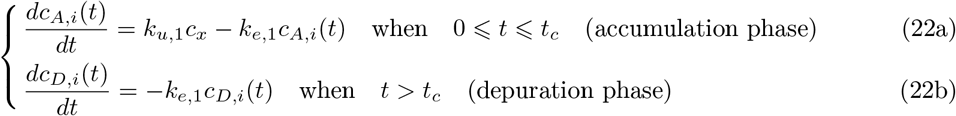

System of equations (22a) and (22b) can easily be solved with the method of the separation of variables (also known as the Fourier method) leading to the following system of solutions for both accumulation and depuration phases:

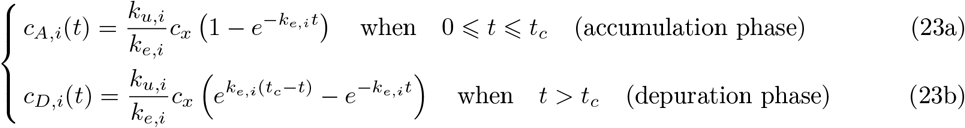

Getting the set of solutions (22) directly from the generic expressions given by the combination of both equations (17) and (21) leads exactly to the same result. Indeed, with one compartment, matrix **E** = –*k_e,i_* and vector **U** = *k_u,i_*.

Inspired from Gestin et al. (2021), when considering only solid black arrows, Figure 3 highlights the target organs that can correspond to one compartment according to *i*: intestine (*i* = 1); cephalon (*i* = 2); caeca (*i* = 3); remaining tissues (*i* = 4). Each compartment has its own parameter pair for uptake (*k_u,i_*) and elimination (*k_e,i_*) rates (Table 2).

**Figure 3:**
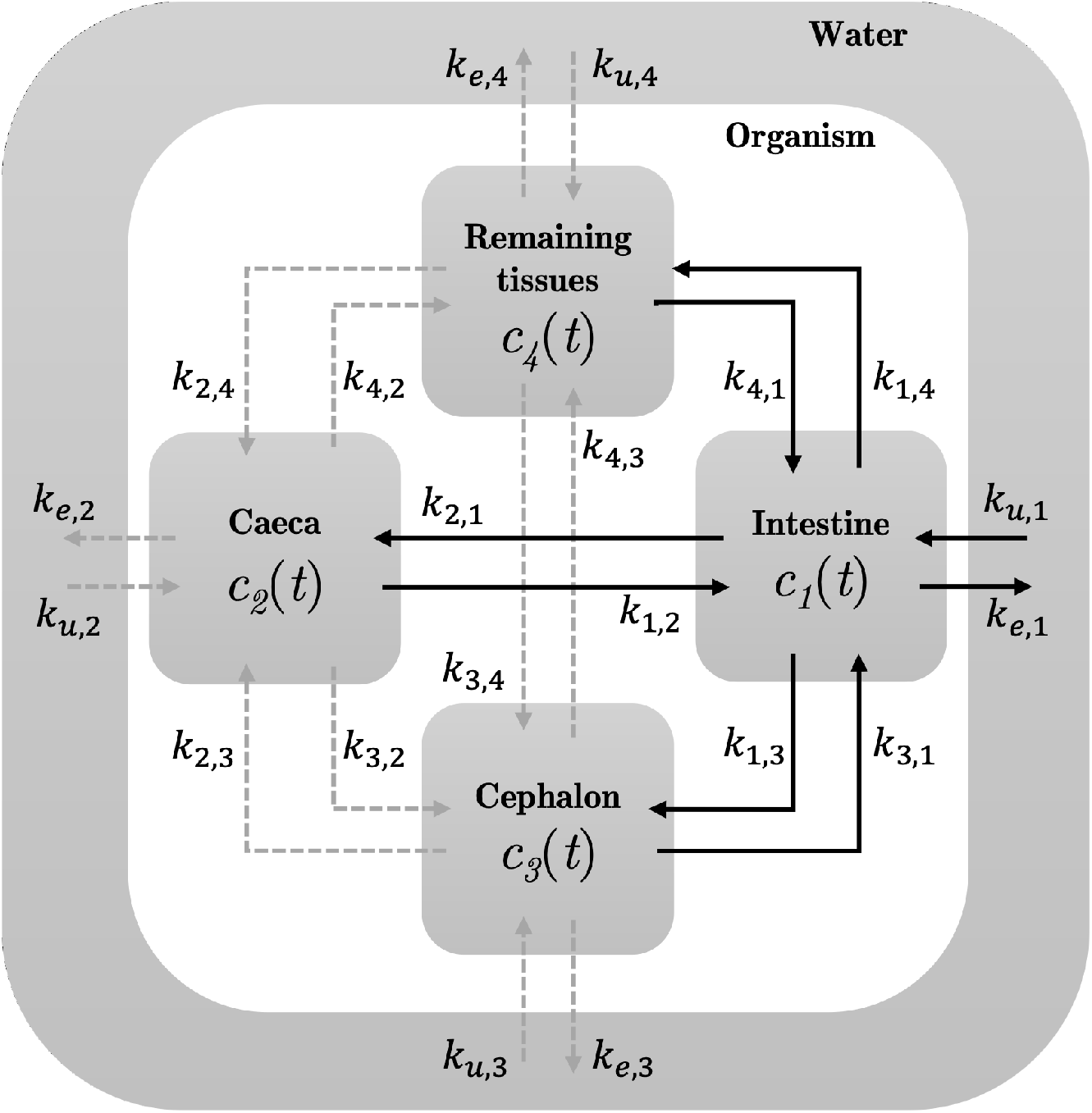
General scheme of the multi-compartment toxicokinetic model that has been used by Gestin et al. (2021) at the initial modelling stage when all compartments were connected to each other. Parameter values as given in Table 3.

**Table 2:** Medians of parameters estimated by Bayesian inference from TK one-compartment models separately fitted to each organ of *Gammarus fossarum* exposed to dissolved Cd at 11.1 *μg.L*^-1^ for 7 days, before being placed for 14 days under depuration conditions.

| Process | Organ | Parameter | Median value (in $[t]^{-1}$) |
| --- | --- | --- | --- |
| Uptake | Intestines | $k_{u,1}$ | 1917 |
| | Caeca | $k_{u,2}$ | 1571 |
| | Cephalons | $k_{u,3}$ | 91.1 |
| | Remaining tissues | $k_{u,4}$ | 135 |
| Elimination | Intestines | $k_{e,1}$ | 0.506 |
| | Caeca | $k_{e,2}$ | 0.053 |
| | Cephalons | $k_{e,3}$ | 0.060 |
| | Remaining tissues | $k_{e,4}$ | 0.026 |

In Gestin et al. (2021), model parameters were estimated under the unified Bayesian framework proposed by Ratier et al. (2019). In particular, parameters of TK one-compartment models were fitted separately for each organ of *G. fossarum* exposed to dissolved Cd at 11.1 *μg.L*^-1^ for 7 days, before being placed for 14 days under depuration conditions. Getting median parameter values as given in Table 2 allows to simulate what happens within intestines when it is connected to all other organs, for example (see SI for more details).

### 4.2 Case study with four compartments

Applying the general matrix ODE system given by the set of equations (15) and (16) to the particular case of four compartments connected by pairs (Figure 3) leads to the following writing:

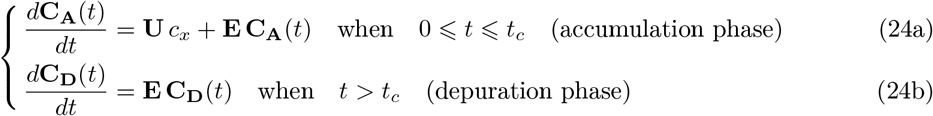

with vectors and matrices defined as follows:

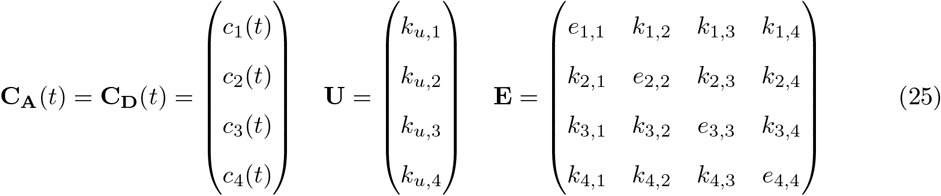

and diagonal elements of matrix **E** defined by:

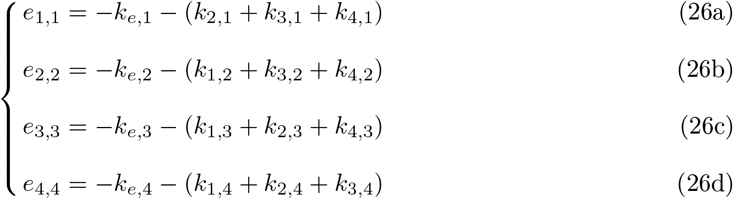

The exact solution of the matrix ODE system (24) can be directly deduced from equations (17) and (21):

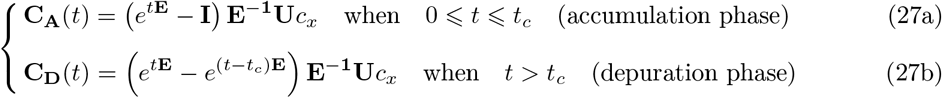

Developing the matrix equations (27a) and (27b) finally provides both following sets of four equations for each accumulation and depuration phases:

- For the accumulation phase (0 ≤ *t* ≤ *t_c_*):

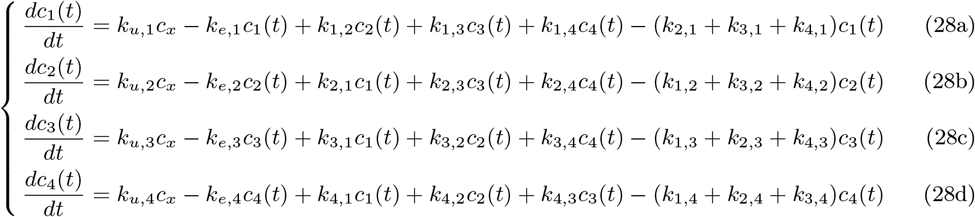
- For the depuration phase (*t > t_c_*):

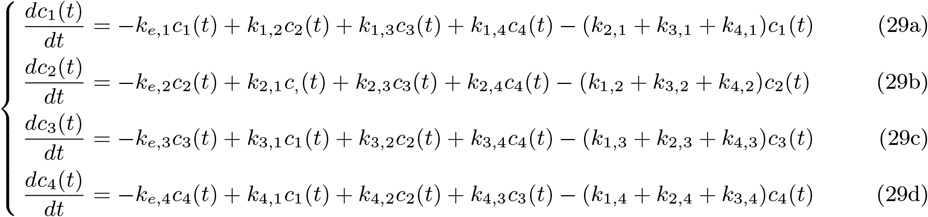

The four compartment model in equations (27a) and (27b), based on the matrix ODE system in equations (24a) and (24b), with its exact solution in (28) and (29), assumes that all compartments are connected to each others by pairs and with the external medium (Figure 3). This means that the model is considering all incoming and outgoing arrows from all compartments. This model comprises a total of 20 parameters, plus the *c_x_* value for the exposure concentration. Median parameter values as given in Table 3 were used to simulate what happens within each organ in terms of internal concentration dynamic, and compared to the previous results with the four independent one-compartment TK models. See SI for details.

**Table 3:** Parameter estimates (expressed as medians and 95% uncertainty intervals) of the four-compartment model corresponding to solid black arrows in Figure 3 as provided by Gestin et al. (2021) in their Table S6. The first column stands for connected organs, either to water or to the other organs; the second column is for parameter names; the next three columns are for medians, lower and upper quantiles of parameter estimates when *G. fossarum* was exposed to Cd = 11.1 *μg.L*^-1^.

| Organ-Connection | Parameter | Median | Q<sub>2.5%</sub> | Q<sub>97.5%</sub> |
| --- | --- | --- | --- | --- |
| Intestines (uptake) | $k_{u,1}$ | 3342 | 2720 | 3707 |
| Intestines (elimination) | $k_{e,1}$ | 0.54 | 0.415 | 1.402 |
| Intestines-Caeca | $k_{21}$ | 0.873 | 0.603 | 1.739 |
| Caeca-Intestines | $k_{12}$ | 0.218 | 0.132 | 0.376 |
| Intestines-Cephalons | $k_{31}$ | 0.059 | 0.034 | 0.124 |
| Cephalons-Intestines | $k_{13}$ | 0.262 | 0.124 | 0.871 |
| Intestines-remaining tissues | $k_{41}$ | 0.069 | 0.049 | 0.126 |
| Remaining tissues-Intestines | $k_{14}$ | 0.14 | 0.086 | 0.238 |
| Intestines | $\sigma_1$ | 8.974 | 6.469 | 15.28 |
| Caeca | $\sigma_2$ | 17.94 | 13.07 | 26.84 |
| Cephalons | $\sigma_3$ | 1.223 | 0.863 | 1.818 |
| Remaining tissues | $\sigma_4$ | 1.468 | 1.06 | 2.242 |

As illustrated above, our generic modelling framework allows to simulate complex situations involving several compartments whatever their connections in pairs and/or with the exposure media. Let’s now relate what we did in subsection 4.1 - that is simulations of four one-compartment models for each organ of *G. fossarum* exposed to Cd - with the four-compartment model developed by Gestin et al. (2021). Indeed, Gestin et al. (2021) showed that *G. fossarum* takes up and eliminates Cd rather quickly, with the intestines and the caeca accumulating and depurating the most compared to the cephalon and the remaining tissues. Gestin et al. (2021) also proved that a four-compartment model better describes the Cd internal contamination route than single one-compartment models for each organ. And they finally highlighted that the most parsimonious multi-compartments model corresponds to the solid black arrows on Figure 3.

Such a situation in fact correspond to a nested model within the four-compartment ODE system as given by equations (24a) and (24b). This model thus comprises only 12 parameters whose values are listed in Table 3. Figure 4 shows the simulated kinetics within the four organs. Our four curves exactly superimpose to the four median curves provided in Figure 3 by Gestin et al. (2021). Benefiting from this exact match between our exact generic solution and what was numerically integrated by Gestin et al. (2021) before the implementation of their inference process ultimately strengthens both approaches: the numerical integration (a basic Euler integration scheme with a time-step equal to 1/10 day); and the curve plotting from the exact solution. Nevertheless, in the perspective to infer parameter values from observed data, there is absolutely no doubt that the exact solution will provide much better computational performance for the implementation of the Monte Carlo Markov Chain simulations needed to use Bayesian inference. Readers who would like to convince themselves of this added value of our generic solving can refer to our preliminary results on the following GitHub repository https://gitlab.in2p3.fr/sandrine.charles/rpbtk. Indeed, based on preliminary results (not shown) embedded within a dedicated R-package (named rPBTK), we already experienced that using this generic exact solution divided at least by 100 the computation time of the whole process. Our first inference implementation required to numerically integrate the original ODE system, then to run the MCMC algorithm to fit the numerical solution to the accumulation-depuration data sets corresponding to the different organs and tissues. The numerical integration step was highly computationally demanding. Avoiding this numerical step, much faster now deliver relevant and precise parameter estimates.

**Figure 4:**
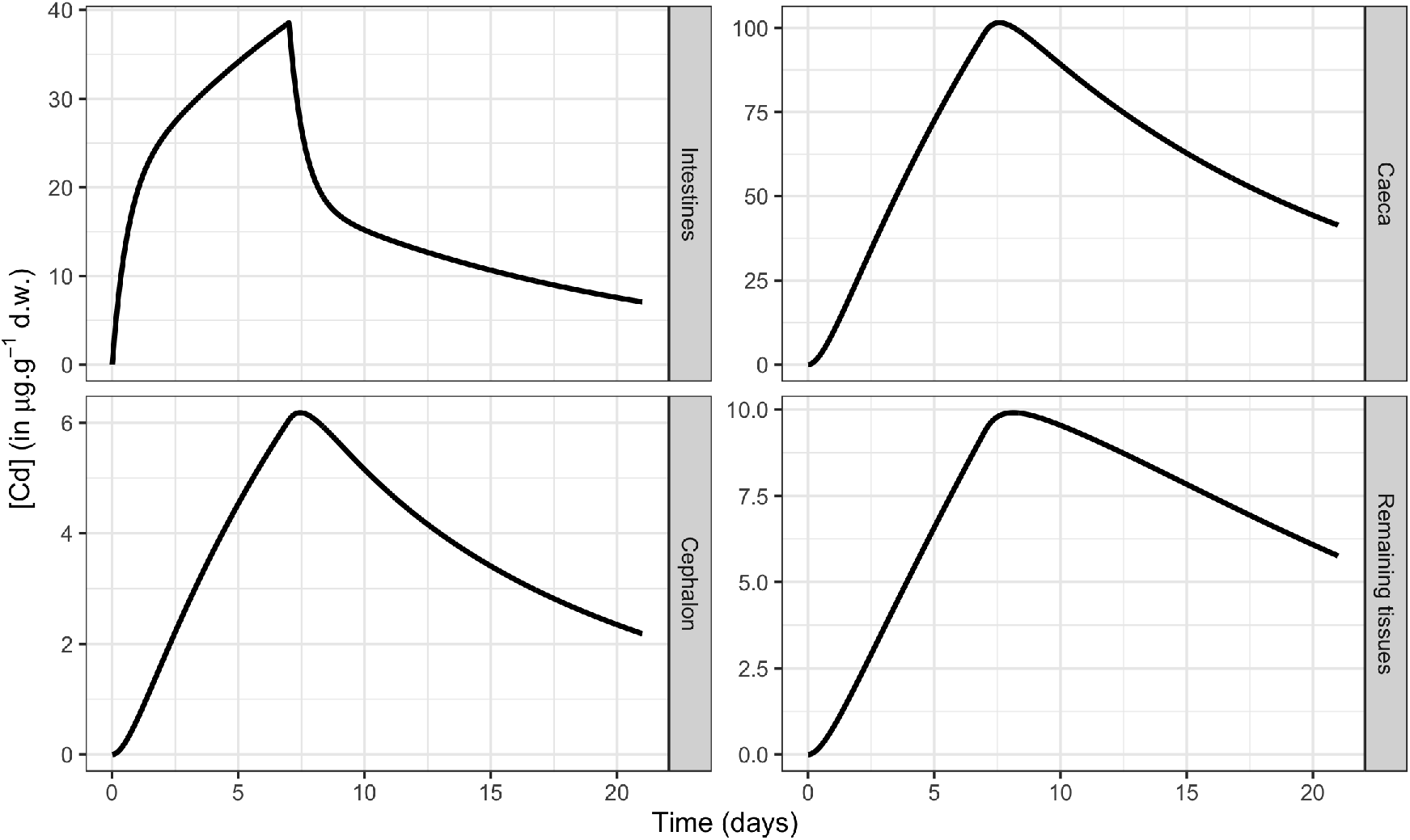
Simulations of the internal concentrations within the different organs or tissues of *Gammarus fossarum* when exposed to an external concentration of cadmium equal to 11.1 *μg.L*^-1^. The four-compartment PBK solution used for simulations is given in equation (27). Parameter values are those given in Table 2.

### 4.3 Case study with six compartments

Here is a final illustration of the usefulness of our generic solution to simulate any PBK model. We identically reproduced all the simulation provided by Zhang et al. (2022), concerning an exposure of zebrafish (*Danio rerio*) to arsenic (As). To this end, Zhang et al. (2022) proposed a six-compartments model with five compartments corresponding to organs: gills (*i =* 2), intestine (*i =* 3), liver (i = 4), head (i = 5) and carcass (i = 6). The sixth compartment corresponds to blood (i = 1). The five organs were connected to blood, while gills and intestine were also connected to the external medium (As contaminated water). These assumptions are translated in Figure 5, together with the different parameters used for the corresponding PBK model (Table 4). From the generic matrix form of a PBK model we present in this paper, the model proposed by Zhang et al. (2022) writes as follows:

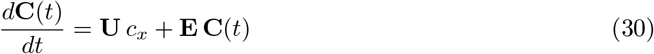

with vectors and matrices defined as follows:

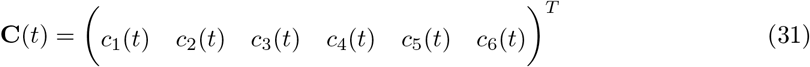

where *c_i_*(*t*), ∀*i* = 1, 6, are the variables corresponding to internal concentrations to be simulated. Variables *c_i_*(*t*) are equal to 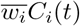, ∀*i* = 1, 6, with 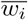 the mean wet weight of blood volume (when *i* = 1) or fish organs (∀*i* = 2, 6). Variables *C_i_*(*t*) are the measured concentrations of As in fish organs (∀*i* = 2, 6, expressed in *μg.g*^-1^) at time *t* (in days). Variable *c_x_* is the exposure concentration of As in water (expressed in *μg.L*^-1^), assumed to be constant over time.

**Figure 5:**
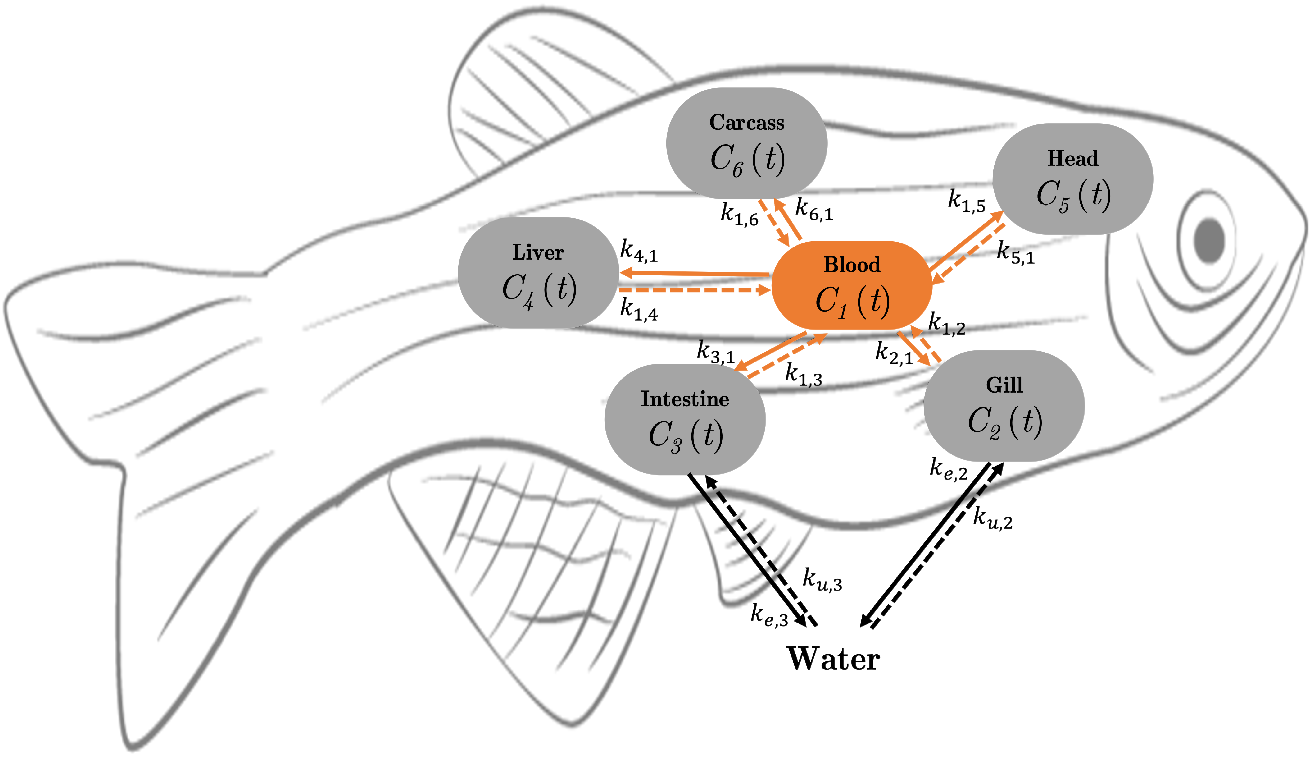
General scheme of the six-compartment kinetic model for zebrafish exposed to arsenic (As, adapted from Zhang et al. (2022)). Parameters, between-organ and/or with-water connections, numerical values and units are given in Table 4.

**Table 4:** Parameter estimates for arsenic distribution in zebrafish (from Zhang et al. (2022)). Notation 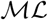 stands for the estimated maximum likelihood value of the parameter.

| Connection | $\mathcal{ML}$ | Value | Unit | Connection | $\mathcal{ML}$ | Value | Unit |
| --- | --- | --- | --- | --- | --- | --- | --- |
| Water to gills | $k_{u,2}$ | $5.28 \cdot 10^{-5}$ | L.d<sup>-1</sup> | Gill to water | $k_{e,2}$ | 0.152 | d<sup>-1</sup> |
| Water to intestine | $k_{u,2}$ | $1.52 \cdot 10^{-4}$ | L.d<sup>-1</sup> | Intestine to water | $k_{e,2}$ | 0.672 | d<sup>-1</sup> |
| Blood to gills | $k_{2,1}$ | 76.0 | d<sup>-1</sup> | Gill to blood | $k_{1,2}$ | 58.2 | d<sup>-1</sup> |
| Blood to intestine | $k_{3,1}$ | 27.8 | d<sup>-1</sup> | Intestine to blood | $k_{1,3}$ | 3.94 | d<sup>-1</sup> |
| Blood to liver | $k_{4,1}$ | 38.5 | d<sup>-1</sup> | Liver to blood | $k_{1,4}$ | 23.6 | d<sup>-1</sup> |
| Blood to head | $k_{5,1}$ | 97.5 | d<sup>-1</sup> | head to blood | $k_{1,5}$ | 8.48 | d<sup>-1</sup> |
| Blood to carcass | $k_{6,1}$ | 7.20 | d<sup>-1</sup> | Carcass to blood | $k_{1,6}$ | 0.317 | d<sup>-1</sup> |

From concentrations within organs, the concentration for the whole organism can be deduced as follows:

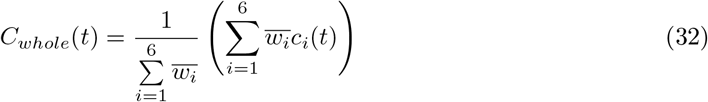

According to our mathematical formalism, uptake parameters from water are included in the following vector:

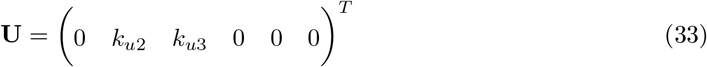

while elimination parameters towards water, as well as parameters corresponding to organ-organ connections, are gathered together within the following matrix:

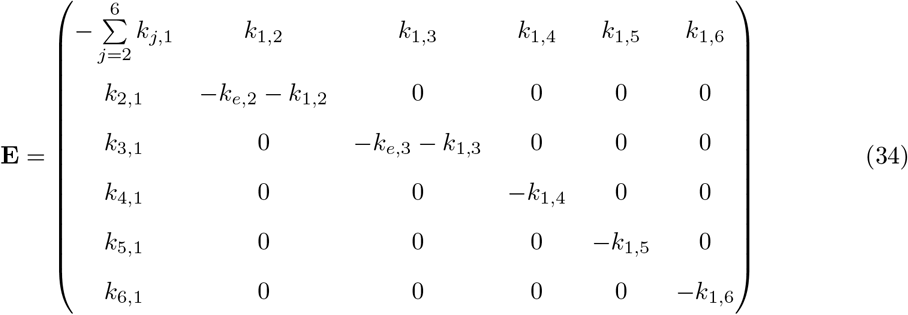

The matrix ODE system (30) finally leads to the following exact solution:

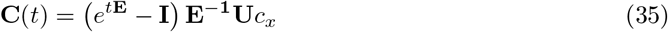

where *c_x_* is the exposure concentration in water.

The above matrix solution (35) can then be developed in order to retrieve the six-compartment PBPK model as constructed by Zhang et al. (2022). Denoting the exposure concentration in water (variable *c_x_* in our modelling) by *C_water_*, we ultimately get:

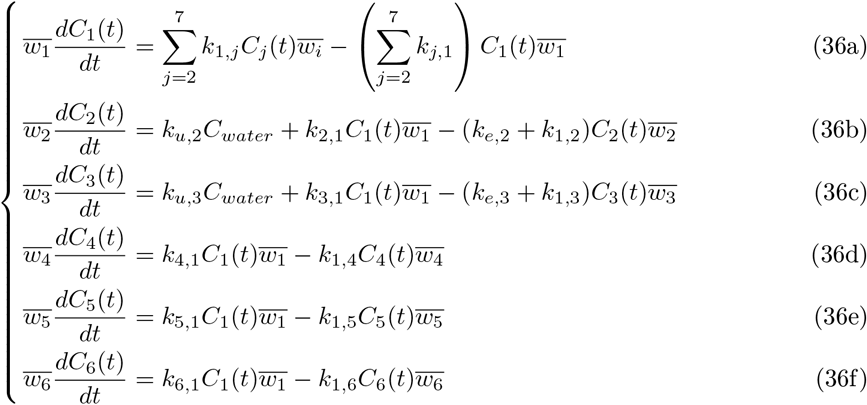

Zhang et al. (2022) measured internal concentrations over time in each organs and in blood during both accumulation and depuration phases. This data allowed them to estimates all their parameters. The use of a Bayesian inference framework also provided them with the quantification of the uncertainty on these parameters. Zhang et al. (2022) then deliver parameter estimates as maximum likelihood (ML) values associated with a 95% credible interval, as partly reported in Table 4.

Based on parameter 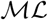 values in Table 4, using fresh weight of the different organs as provided in Table S2 from Zhang et al. (2022), we performed simulations of the internal concentrations within each of the five organs as well as in blood (variables *c_i_*(*t*), ∀*i* = 1, 6). As shown on Figure 6, our generic solved TK model (see equation (35)) again exactly reproduce the median curves provided by Zhang et al. (2022) in their Figure 1 (solid blue lines). Such an exact match between our curves and the authors’ ones again provide a full check of the mathematical writing of our generic solution, together with the fact that it produces identical median curves. These results again reinforce the possibility of using this exact generic solution for the further implementation of inference processes with the guarantee of obtaining the parameter estimates much faster, avoiding the time-consuming step of the numerical integration of the ODE system.

**Figure 6:**
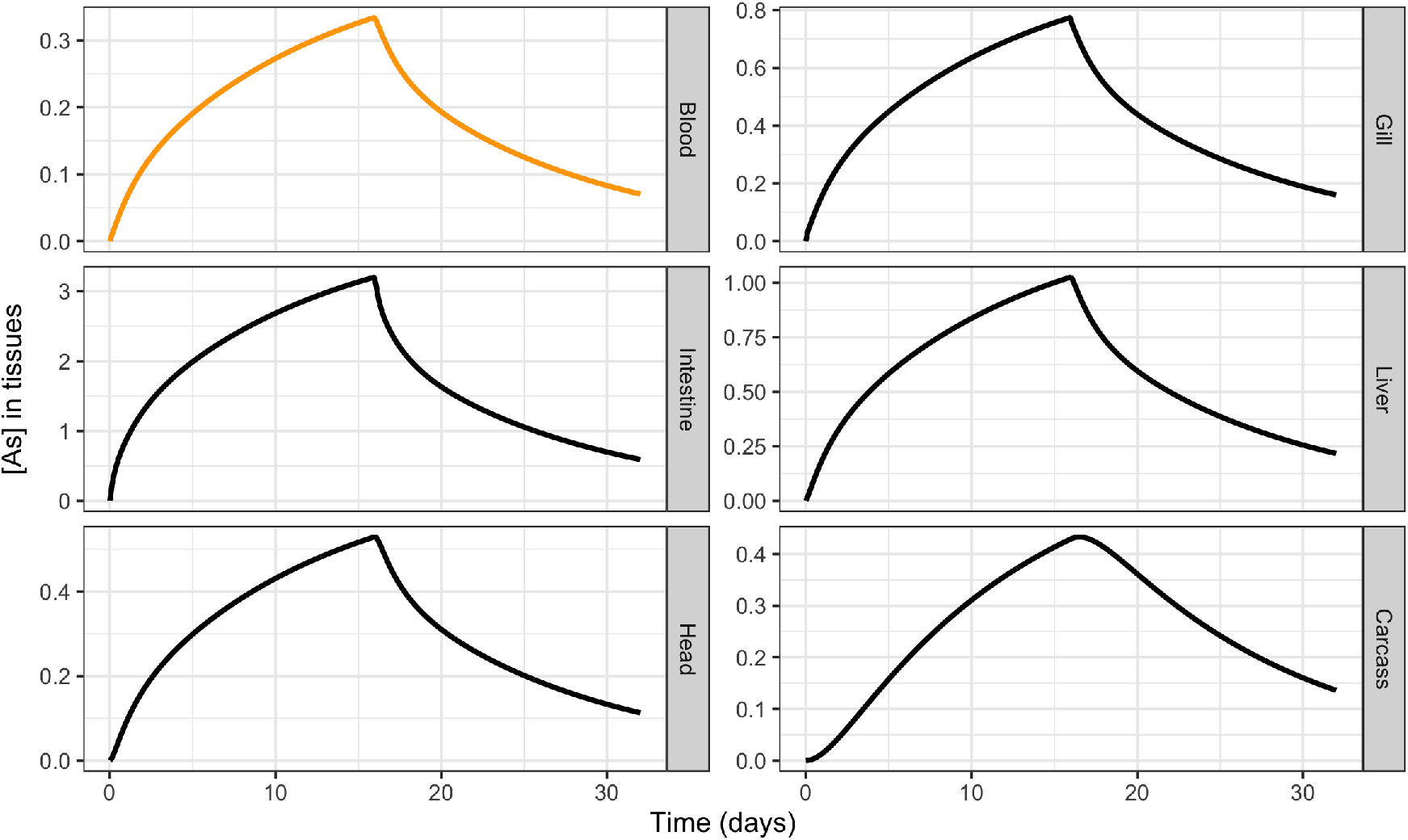
Simulation of the total arsenic concentration in fish tissues and blood during the accumulation phase of 16 days (exposure concentration equal to 400 *μg.L*^-1^) and the subsequent 16 days of depuration in clean medium. The PBK model used for simulations is given as a matrix solution in equation (35) with parameter values from Table 4.

## 5 Conclusion

Our generic solving of any full PBK model comprising as many compartments as physiologically needed, as well as all potential connections between compartments and with the external medium, revealed particularly efficient in simulating diverse situations in terms of species, compounds and purpose. The four-compartment PBK model for *G. fossarum* exposed to Cd highlighted the dynamic transfer of Cd among the different organs. The six-compartment PBK model for *D. rerio* exposed to As brought to light that intestines were the main uptake site for waterborne As, instead of gills as authors’ expected. Several other examples have complementary been tested (results not shown). Nevertheless, the genericity of the solution we proposed here could still be further extended in order to account for simultaneous but different routes of exposure (via water, food, sediment and/or pore water for example), as well as several elimination processes, among which the dilution by growth would allows to be more realistic for long-lived species. Moreover, account for the metabolization of the parent compound, at least in phase-I metabolites, could also be of great interest to better deal with organic compounds.

Benefiting from this generic solution also allows us to envisage the continuation of this work with confidence, in particular the implementation of a Bayesian inference framework to get parameter estimates in a quick and efficient way, based on a fully harmonized methodology. The next step will thus be to make freely and easily accessible this generic modelling framework, innovative both in the writing of the PBK model and in its exact resolution, but also in the implementation of the fitting and simulation tools. Most users, who are not necessarily modelling specialists, would be willing to use more complex PBK models, especially if necessity dictates and sufficient input data is available. Unfortunately, turnkey tools are lacking. In line with the MOSAIC platform spirit, we have therefore already undertaken the creation of a new online service that will facilitate the use of PBK models, also for beginners, with a step-by-step workflow to first upload data and select the model to fit. By default, the most complete one will be automatically proposed from which users will be able to deselect either compartments or exchanges between compartment and/or with water according to prior physiological knowledge or to try nested models and finally identify the most appropriate. Then, users will be accompanied to run the fitting process, get the results (parameter estimates and fitting plots), look at the goodness-of-fit criteria (with guidance on their interpretation), and use the model comparison criteria in case several models would have been tested. Once this calibration step achieved, users will have the possibility to run simulations, to compare with additional data or to plan further experiments for example. In the end, all expected features of a convenient help-decision service could be offered, with a particular attention in further supporting the next generation tiered PBK modeling framework, that could become the new paradigm in human and environmental risk assessment. Indeed, once our generic PBK modelling framework firmly anchored in practice, we would be in the right position to consider its coupling with mechanistic models to build a completely general modelling framework from exposure (pharmaco/toxico-kinetics) to effects (pharmaco/toxico-dynamics) on life history traits, hence defining a unifying PBKD modelling framework.

## Acknowledgments

The authors would like to thank the BioEEnViS research federation (Biodiversity, Water, Environment, City, Health) for its financial support allowing to collaboratively achieve this study. Moreover, this work is part of the ANR project APPROve (ANR-18-CE34-0013) for an integrated approach to propose proteomics for biomonitoring: accumulation, fate and multi-markers (https://anr.fr/Projet-ANR-18-CE34-0013). A large part of the work also benefited from the French GDR “Aquatic Ecotoxicology” framework which aims at fostering stimulating scientific discussions and collaborations for more integrative approaches. The authors would like to express their sincere gratitude to Julie KLEINE-SCHULTJANN as part of a training course during her 5^th^ year study at the National Institute of Applied Sciences (INSA) in Lyon (France). At last, the work presented in this paper was performed using the computing facilities of the CC LBBE/PRABI.

## Author contribution

SC resolved the full generic PBK model, wrote the manuscript an coordinated all discussion around this paper. OGes, AC, OGef, TLL and CL were strongly involved in the *G. fossarum* case study, in a close collaboration within OGes’s PhD supervision, who led all the underlying experimental work. JB, DL and VB took care with special attention with the model equations writing. VB wrote the R source code which generated simulation curves; this R code was also checked by SC, JB, DL and CL. All authors contributed to and agreed on the final version of the manuscript.

## Supplementary information

Supplementation information is available at our dedicated Zenodo repository: https://zenodo.org/record/6501782. more information and details may be asked directly to the corresponding author.

