## Supplementary Information for "Generic solving of physiologically-based kinetic models in support of next generation risk assessment due to chemicals"

Sandrine CHARLES*et al.

### Supplementary Information

May 16, 2022

---

**References** **14**

### 1 Details on the generic solving of the PBK model

This supplementary material contains all intermediate details to get the generic solutions presented in the manuscript in their final form. The starting point is the full matrix ODE system with $n$ compartments all related by pairs writing as follows:

$$\frac{d\mathbf{C}(t)}{dt} = \mathbf{U}\,c_x + \mathbf{E}\,\mathbf{C}(t) \tag{1}$$

which has the following final particular solution corresponding to the initial condition $\mathbf{C_{wsm}}(t = 0) = \mathbf{C_0}$, where index **wsm** stands for with second member.:

$$\mathbf{C_{wsm}}(t) = \left(\int_0^t e^{(t-\tau)\mathbf{E}} d\tau\right) \mathbf{U} c_x + e^{t\mathbf{E}}\mathbf{C_0} \tag{2}$$

#### 1.1 Generic solving based on the use of matrix exponential

In this subsection, we detail how to simplify a matrix integral from the definition of a matrix exponential. The aim is thus to get a simplified expression of:

$$\int_0^t e^{(t-\tau)\mathbf{E}} d\tau \tag{3}$$

From the general expression of a matrix exponential, we get:

$$e^{(t-\tau)\mathbf{E}} = \sum_{k=0}^{\infty} \frac{1}{k!} \left((t-\tau)\mathbf{E}\right)^k \tag{4}$$

Hence:

$$
\begin{aligned}
\int_0^t e^{(t-\tau)\mathbf{E}} d\tau &= \int_0^t \left( \sum_{k=0}^{\infty} \frac{1}{k!} \left( (t-\tau)\mathbf{E} \right)^k \right) d\tau \\
&= \sum_{k=0}^{\infty} \left( \frac{\mathbf{E}^k}{k!} \int_0^t (t-\tau)^k d\tau \right) = \sum_{k=0}^{\infty} \left( \frac{\mathbf{E}^k}{k!} \Big[ - \frac{(t-\tau)^{k+1}}{k+1} \Big]_0^t \right) \\
&= \sum_{k=0}^{\infty} \left( \frac{\mathbf{E}^k}{k!} \left( -0 + \frac{t^{k+1}}{k+1} \right) \right) = \sum_{k=0}^{\infty} \left( \frac{t^{k+1}\mathbf{E}^k}{(k+1)!} \mathbf{E}\mathbf{E}^{\mathbf{-1}} \right) \\
&= \sum_{k=0}^{\infty} \frac{t^{k+1}\mathbf{E}^{k+1}}{(k+1)!} \mathbf{E}^{\mathbf{-1}}
\end{aligned}
$$

Let's denote $\ell = k + 1$. From the above expression we thus get:

$$
\begin{aligned}
\int_0^t e^{(t-\tau)\mathbf{E}} d\tau &= \sum_{\ell=1}^{\infty} \frac{t^\ell \mathbf{E}^\ell}{\ell!} \mathbf{E}^{\mathbf{-1}} \\
&= \sum_{\ell=0}^{\infty} \left( \frac{t^\ell \mathbf{E}^\ell}{\ell!} - \frac{t^0 \mathbf{E}^0}{0!} \right) \mathbf{E}^{\mathbf{-1}} = \sum_{\ell=0}^{\infty} \left( \frac{t^\ell \mathbf{E}^\ell}{\ell!} - \mathbf{I} \right) \mathbf{E}^{\mathbf{-1}} \\
&= \left( e^{t\mathbf{E}} - \mathbf{I} \right) \mathbf{E}^{\mathbf{-1}}
\end{aligned}
$$

We finally obtain the following matrix final solution:

$$
\mathbf{C_{wsm}}(t) = \left( e^{t\mathbf{E}} - \mathbf{I} \right) \mathbf{E}^{\mathbf{-1}} \mathbf{U} c_x + e^{t\mathbf{E}} \mathbf{C_0} \tag{7}
$$

#### 1.2 Generic solving based on Jordan normal forms

Mathematically speaking, any square matrix is similar to its Jordan normal form $\mathbf{J}$ via an appropriate transition matrix $\mathbf{P}$. Hence, any matrix $\mathbf{E}$ can be written as follows:

$$
\mathbf{E} = \mathbf{P}\mathbf{J}\mathbf{P}^{\mathbf{-1}} \tag{8}
$$

where matrix $\mathbf{P}$ is the transition matrix defined with columns equal to eigenvectors of matrix $\mathbf{E}$. Because eigenvectors are forming a base, matrix $\mathbf{P}$ is always invertible.

Considering real Jordan normal forms, matrix $\mathbf{J}$ will be a block diagonal matrix formed of real Jordan blocks, that are themselves real matrix blocks composed of zeroes everywhere except on the diagonal, filled with fixed elements $\lambda_i \in \mathbb{R}$ $(i = \overline{1, n})$, and the upper-diagonal filled with ones. Elements $\lambda_i$ $(i = \overline{1, n})$ correspond to the $n$ eigenvalues associated with the $n$ eigenvectors of matrix $\mathbf{E}$ as used to build matrix $\mathbf{P}$.

Then, it immediately comes that:

$$e^{t\mathbf{E}} = \mathbf{P}\mathbf{e}^{\mathbf{tJ}}\mathbf{P}^{\mathbf{-1}} \tag{9}$$

Equivalently, we also get the following expressions:

$$e^{(t-\tau)\mathbf{E}} = \mathbf{P}e^{(t-\tau)\mathbf{J}}\mathbf{P}^{\mathbf{-1}} \tag{10}$$

and

$$\int_0^t e^{(t-\tau)\mathbf{E}} d\tau = \mathbf{P}\left(\int_0^t e^{(t-\tau)\mathbf{J}} d\tau\right)\mathbf{P}^{\mathbf{-1}} \tag{11}$$

Considering the set of complex values $\mathbb{C}$, the Jordan normal form can be simply written as a diagonal matrix:

$$\mathbf{J} = diag\left\{\lambda_i\right\}_{i=1,n} \quad \text{with} \quad \lambda_i \in \mathbb{C} \tag{12}$$

leading to

$$e^{(t-\tau)\mathbf{J}} = diag\left\{e^{(t-\tau)\lambda_i}\right\}_{i=1,n} \tag{13}$$

For each compartment $i$ ($i = 1, n$), we can then calculate:

$$\int_0^t e^{(t-\tau)\lambda_i} d\tau = \frac{1}{\lambda_i}\left(e^{\lambda_i t} - 1\right) \quad \forall i = 1, n \tag{14}$$

what finally leads to the following writing:

$$\int_0^t e^{(t-\tau)\mathbf{J}} d\tau = diag\left\{\frac{1}{\lambda_i}\left(e^{\lambda_i t} - 1\right)\right\}_{i=1,n} \tag{15}$$

then to the following expression:

$$\int_0^t e^{(t-\tau)\mathbf{E}} d\tau = \mathbf{P}\, diag\left\{\frac{1}{\lambda_i}\left(e^{\lambda_i t} - 1\right)\right\}_{i=1,n}\mathbf{P}^{\mathbf{-1}} \tag{16}$$

This finally leads to the following final exact solution:

$$\mathbf{C_{wsm}}(t) = \mathbf{P}\, diag \left\{ \frac{1}{\lambda_i} \left( e^{\lambda_i t} - 1 \right) \right\}_{i=1,n} \mathbf{P^{-1}}\, \mathbf{U}\, c_x + e^{t\mathbf{E}}\, \mathbf{C_0} \tag{17}$$

### 2 R command lines and simulation results

```r
# Clean working space
rm(list = ls())
# Load required packages
library(deSolve)

## Error in library(deSolve):  there is no package called 'deSolve'
```

#### 2.1 One-compartment PBK model

Below are R command lines to simulate the four one-compartment PBK models as done by [1].

```r
# Write the one-compartment TK model for both phases (maccu, mdepu)
# as internal concentrations over time for each phase (caccu, cdepu)
# x stands for time as required by the R function `curve()`
# cx stands for the contaminant exposure concentration
# ku stands for the uptake rate
# ke stands for the elimination rate
maccu <- function(x, cx, ku, ke){ # Accumulation phase
    caccu <- (ku * cx / ke) * (1 - exp(- ke * x))
    return(caccu / 1000)
    }
mdepu <- function(x, cx, ku, ke){ # Depuration phase
    cdepu <- (ku * cx / ke) * (exp(ke * (tacc - x)) - exp(- ke * x))
    return(cdepu / 1000)
    }
```

```r
# Simulations for one single compartment
```

```
par(mar = c(4, 5, 0.1, 0.1), mfrow = c(1, 1))
cx <- 11.1 # in micro-gram/liter
tacc <- 7 # accumulation duration (in days)
tfin <- tacc + 14 # total duration
# Parameter values for the intestines
ku <- 1917 # per day
ke <- 0.506 # per day
# Simulate the internal concentration for accumulation
curve(maccu(x, cx, ku, ke), from = 0, to = tacc,
    xlim = c(0, tfin), las = 1, lwd = 2, xlab = "Time (days)",
    ylab = expression(paste("[Cd] (in ",mu,"g.",g-1," d.w.)")))
# Simulate the internal concentration for depuration
curve(mdepu(x, cx, ku, ke), from = tacc, to = tfin,
    lwd = 2, add = TRUE)
# add a vertical line delimiting both phases
abline(v = tacc, lty = 2)
# Add a legend
legend("topright", legend = "(1)", bty = "n")
```

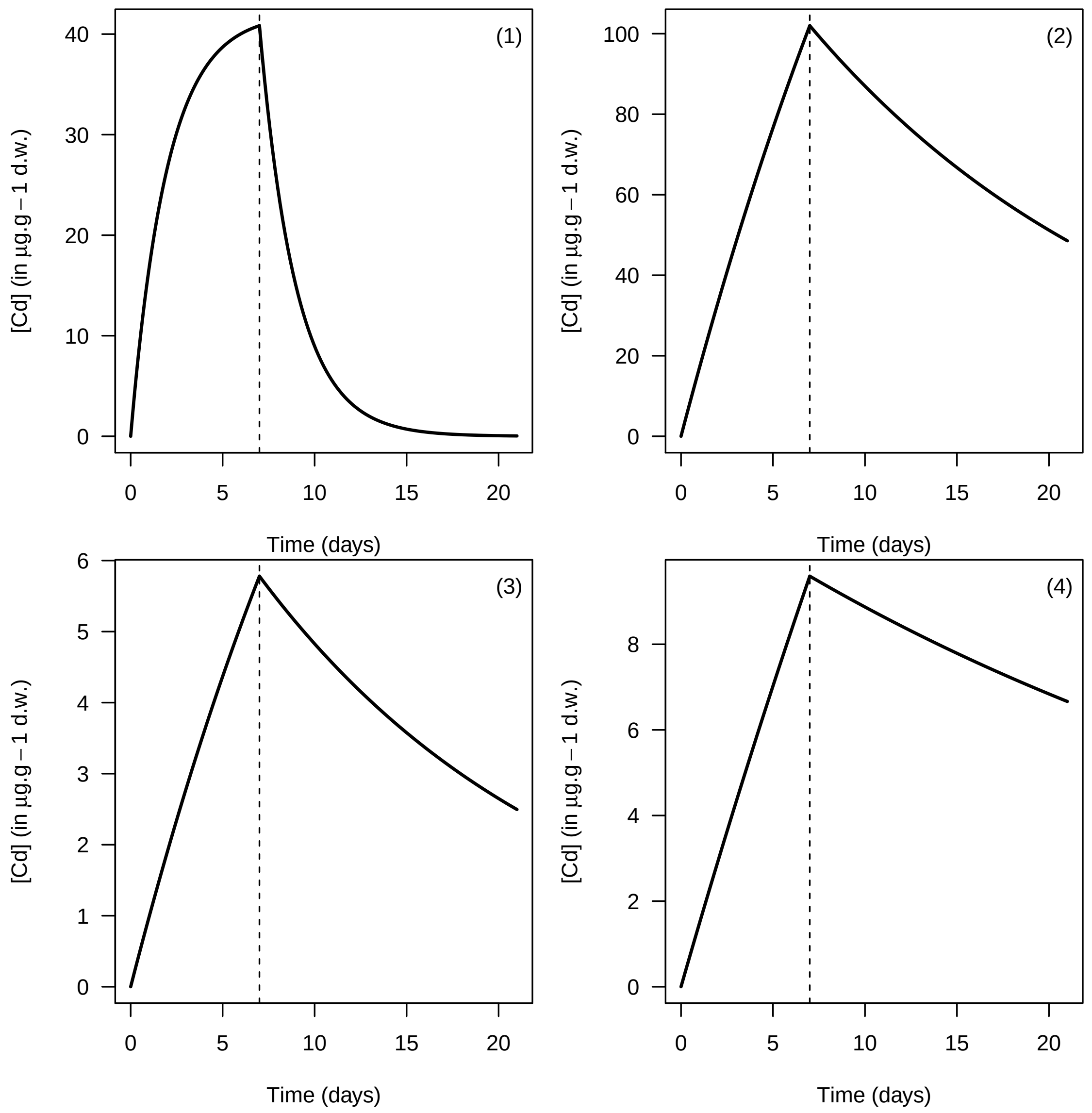

Figure 1: Simulation of bioaccumulation within *Gammarus fossarum* exposed to Cd at concentration 11.1 $\mu g.L^{-1}$. The solid lines stand for simulated internal concentrations; vertical dotted lines delimit accumulation from depuration phases, in (1) intestines, (2) caeca, (3) Cephalons and (4) remaining tissues.

#### 2.2 Four-compartment PBK model

##### 2.2.1 Connecting all compartments by pairs

Always from the study conducted by [1], we compared two types of simulations: (1) simulation from the exact solution as given by both matrix equations (??) and (??); (2) simulations based on the numerical integration of the ODE system of four equations, with the R-package `deSolve` [4], function `ode()` [3, 2].

Parameter estimates used to simulate the four-compartments model when all compartment are connected by pairs are given in Table 1 below. Simulations are only based on medians values.

Table 1: Medians of parameters estimated from the four-compartment TK model simultaneously fitted to each data set corresponding to the four identified organs of *Gammarus fossarum* exposed to dissolved Cd at 11.1 $\mu g.L^{-1}$ for 7 days, before being placed for 14 days under depuration conditions.

| Organ | Parameter | Mean | Median | 2.5% quantile | 97.5% quantile |
|---|---|---|---|---|---|
| Intestines | $k_{u,1}$ | 2600 | 1900 | 0.014 | 13000 |
| Intestines | $k_{e,1}$ | 0.71 | 0.58 | 0.00013 | 3.2 |
| Caeca | $k_{u,2}$ | 1300 | 1600 | 0.00027 | 1800 |
| Caeca | $k_{e,2}$ | 0.0072 | 0.00076 | $1.20.10^{-5}$ | 0.054 |
| Cephalons | $k_{u,3}$ | 190 | 0.16 | $1.60.10^{-5}$ | 2200 |
| Cephalons | $k_{e,3}$ | 0.35 | 0.0089 | $1.40.10^{-5}$ | 3 |
| Remaining tissues | $k_{u,4}$ | 180 | 0.12 | $1.50.10^{-5}$ | 2000 |
| Remaining tissues | $k_{e,4}$ | 0.21 | 0.0041 | $1.40.10^{-5}$ | 2.4 |
| Intestines-Caeca | $k_{2,1}$ | 0.07 | 0.0013 | $1.30.10^{-5}$ | 0.78 |
| Intestines-Caeca | $k_{1,2}$ | 0.033 | 0.017 | $1.50.10^{-5}$ | 0.11 |
| Intestines-Cephalons | $k_{3,1}$ | 0.047 | 0.0025 | $1.30.10^{-5}$ | 0.47 |
| Intestines-Cephalons | $k_{1,3}$ | 0.49 | 0.034 | $1.50.10^{-5}$ | 3.1 |
| Intestines-tissues | $k_{4,1}$ | 0.057 | 0.0035 | $1.30.10^{-5}$ | 0.54 |
| Intestines-tissues | $k_{1,4}$ | 0.34 | 0.032 | $1.50.10^{-5}$ | 2.3 |
| Caeca-Cephalons | $k_{3,2}$ | 0.027 | 0.0037 | $1.40.10^{-5}$ | 0.16 |
| Caeca-Cephalons | $k_{2,3}$ | 0.41 | 0.01 | $1.40.10^{-5}$ | 3.3 |
| Caeca-tissues | $k_{4,2}$ | 0.047 | 0.022 | $1.80.10^{-5}$ | 0.26 |
| Caeca-tissues | $k_{2,4}$ | 0.3 | 0.013 | $1.50.10^{-5}$ | 2.5 |
| Cephalons-tissues | $k_{4,3}$ | 0.35 | 0.013 | $1.40.10^{-5}$ | 2.7 |
| Cephalons-tissues | $k_{3,4}$ | 0.18 | 0.0085 | $1.40.10^{-5}$ | 1.5 |

As a first step, parameter estimates need to be loaded within the R software: The corresponding tabular file, entitled `param4comp.txt`, with parameter estimates provided in the SI on-line within the dedicated repository, available at our dedicated Zenodo repository https://zenodo.org/record/6501782. The script with the R command lines, entitled `script4comp.R`, is also downloadable from this repository.

Figures hereafter illustrate the final results of both simulation outputs, either based on the exact solution (Figure 1), or on the the numerical integration of the ODE system (Figure 2). These figures confirm the exact match between our generic solution of the multi-compartment TK model, with a numerical simulation for given parameter values.

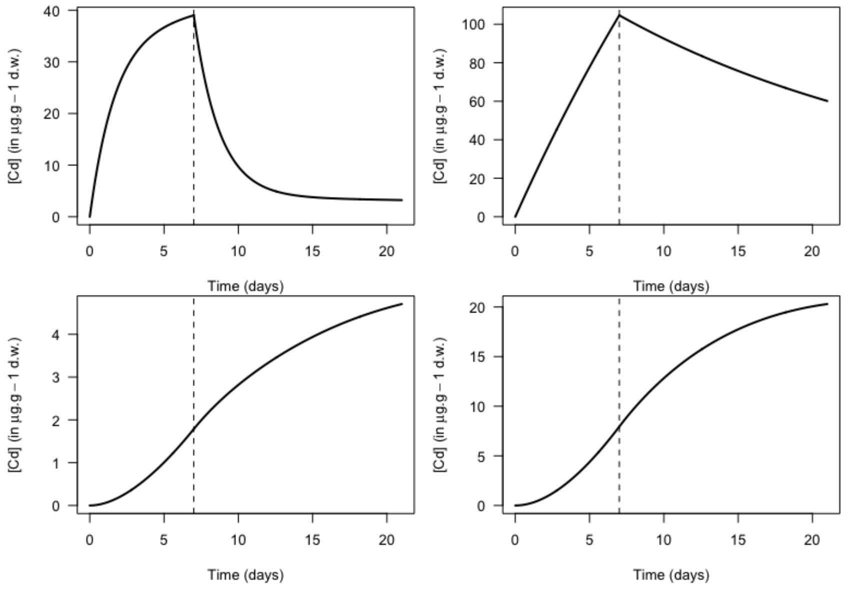

Figure 1: Simulation of the ODE matrix system from its exact solution given by equations (**??**) and (**??**) in all compartments. Parameter values are given in Table 1.

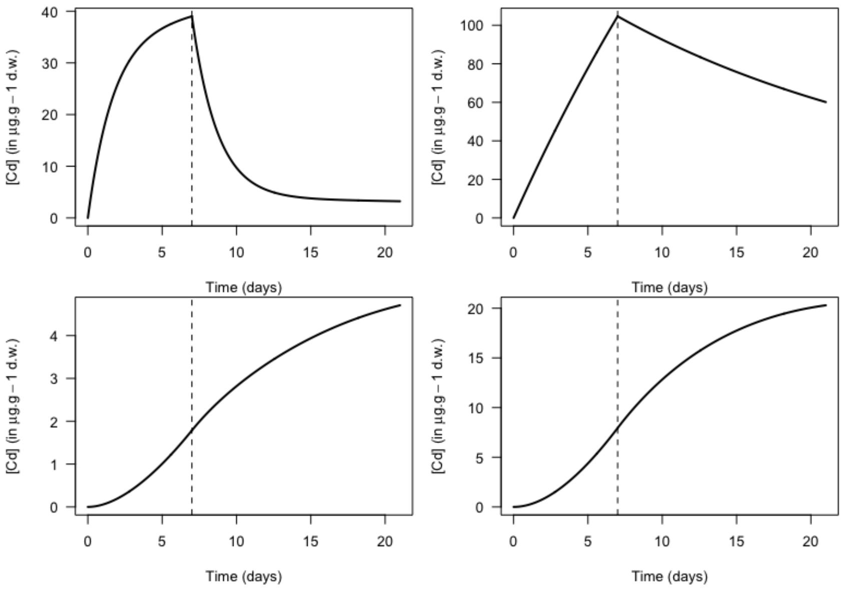

Figure 2: Simulation of the ODE matrix system given in equations (??) and (??), based on a numerical integration. Parameter values are given in Table 1.

##### 2.2.2 Biologically-based connections between compartments

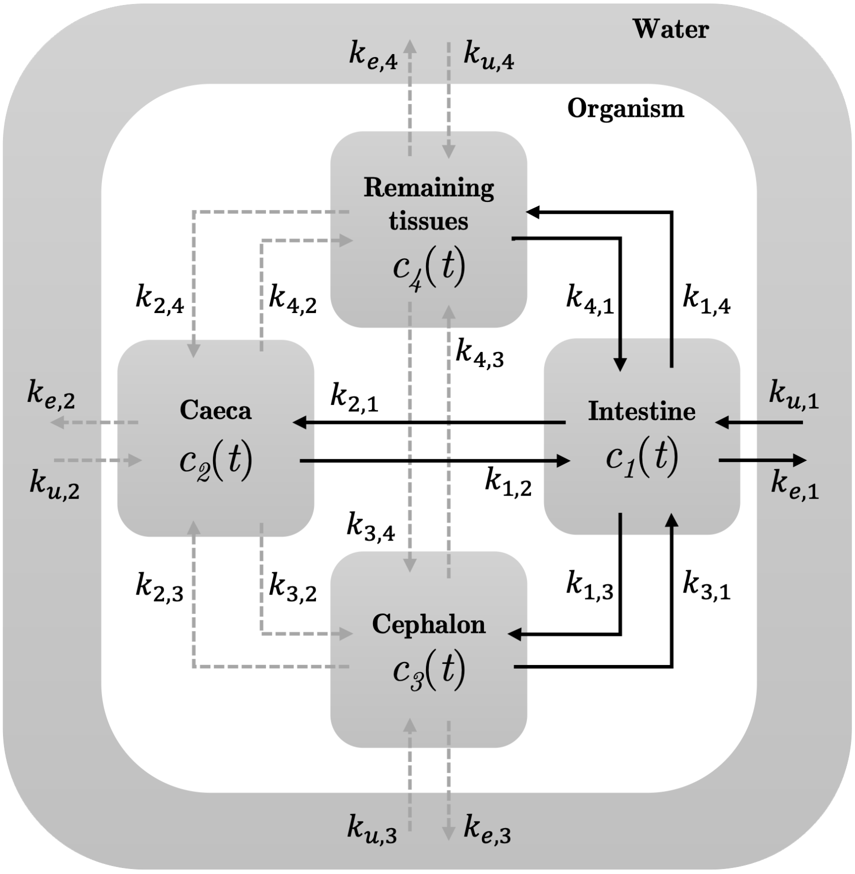

Figure 3: General scheme of the multi-compartment toxicokinetic model that has been used by [1] at the initial modelling stage when all compartments were connected to each other. Parameter values as given in Table 1.

Based on the final results of [1], we provide below two sets of simulations both based on a four-compartment model but only considering biologically-founded compartment connections according to Figure 3 and the black solid arrows. This model assumes that only intestines are directly connected to the external medium (here, water), and accounts for connections only between intestines and the three other organs. The first set of simulations corresponds to internal concentration measured within organs of *G. fossarum* when exposed to Cd at concentration 11.1 $\mu g.L^{-1}$(Figure 4). The second set of simulations is for *G. fossarum* exposed to Hg at concentration 0.27 $\mu g.L^{-1}$ (Figure 5). Parameter estimates are listed in Table 2 for both compounds.

Tabular files with parameters as well as both R scripts are available in the Zenodo repository at `https://zenodo.org/record/6501782`.

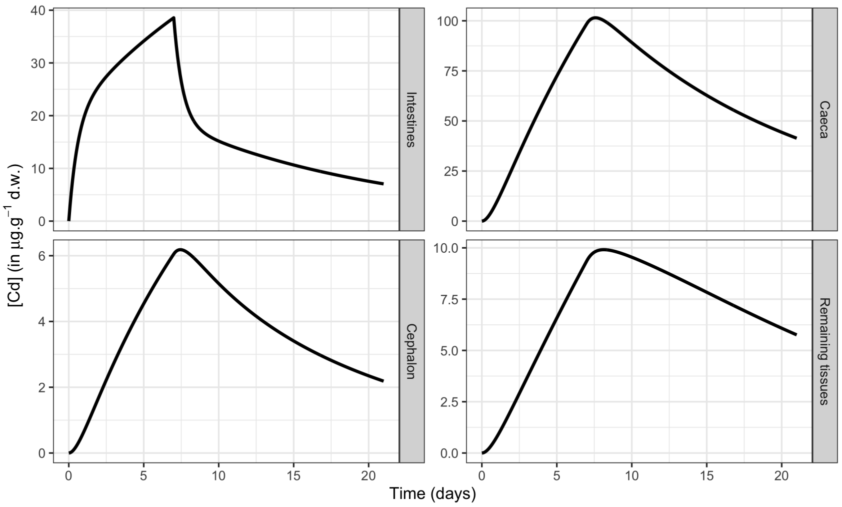

Figure 4: Simulation of the ODE matrix system given in equations (??) and (??), based on a numerical integration. Parameter values are given in Table 2.

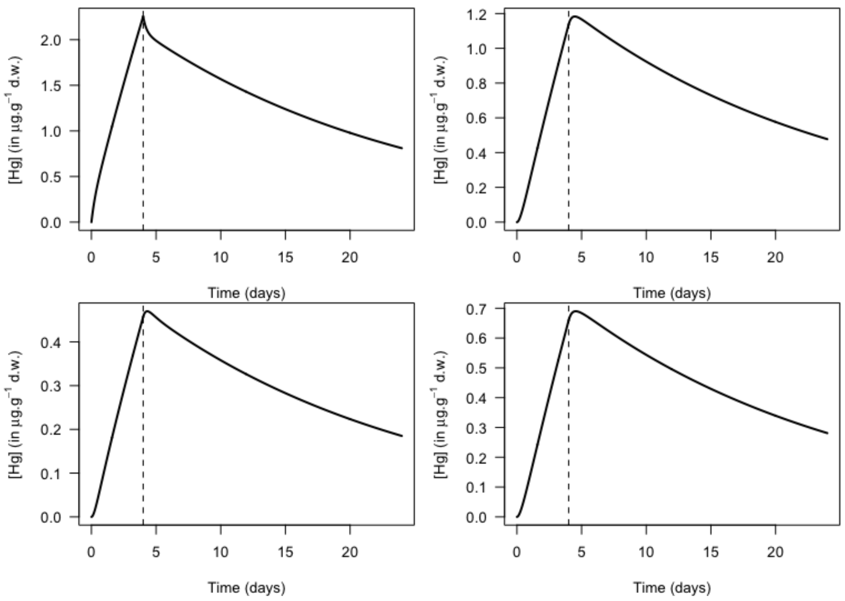

Figure 5: Simulation of the ODE matrix system given in equations (??) and (??), based on a numerical integration. Parameter values are given in Table 2.

| Organ-Connection | Parameter | Median | $Q_{2.5\%}$ | $Q_{97.5\%}$ | Median | $Q_{2.5\%}$ | $Q_{97.5\%}$ |
|---|---|---|---|---|---|---|---|
| Intestines (uptake) | $k_{u,1}$ | 3342 | 2720 | 3707 | 4640 | 3890 | 5272 |
| Intestines (elimination) | $k_{e,1}$ | 0.54 | 0.415 | 1.402 | 0.102 | 0.06 | 0.141 |
| Intestines-Caeca | $k_{21}$ | 0.873 | 0.603 | 1.739 | 1.023 | 0.622 | 1.35 |
| Caeca-Intestines | $k_{12}$ | 0.218 | 0.132 | 0.376 | 1.784 | 0.872 | 2.312 |
| Intestines-Cephalons | $k_{31}$ | 0.059 | 0.034 | 0.124 | 0.515 | 0.269 | 0.653 |
| Cephalons-Intestines | $k_{13}$ | 0.262 | 0.124 | 0.871 | 2.303 | 0.757 | 2.967 |
| Intestines-Residues | $k_{41}$ | 0.069 | 0.049 | 0.126 | 0.552 | 0.405 | 0.714 |
| Residues-Intestines | $k_{14}$ | 0.14 | 0.086 | 0.238 | 1.639 | 0.999 | 2.145 |
| Intestines | $\sigma_1$ | 8.974 | 6.469 | 15.28 | 0.743 | 0.556 | 1.053 |
| Caeca | $\sigma_2$ | 17.94 | 13.07 | 26.84 | 0.434 | 0.323 | 0.615 |
| Cephalons | $\sigma_3$ | 1.223 | 0.863 | 1.818 | 0.076 | 0.056 | 0.113 |
| Residues | $\sigma_4$ | 1.468 | 1.06 | 2.242 | 0.068 | 0.05 | 0.099 |

Table 2: Parameter estimates (expressed as medians and 95% uncertainty intervals) of the four-compartment model corresponding to black arrows in Figure 3 as provided by [1] in their Table S6. The first column stands for connected organs, either to water or to the other organs (see Figure 3, solid black arrows); the second column is for parameter names; the next three columns are for medians, lower and upper quantiles of parameter estimates when *G. fossarum* were exposed to Cd $= 11.1\ \mu g.L^{-1}$; the last three columns are for medians, lower and upper quantiles of parameter estimates when *G. fossarum* were exposed to Hg $= 0.27\ \mu g.L^{-1}$.

##### 2.3 Six-compartment PBK model

This example was build from the work of [5]. You will find the corresponding R script, entitled `script6comp-zhang.R`, within our Zenodo repository at `https://zenodo.org/record/6501782`.

#### References

[1] O. Gestin, T. Lacoue-labarthe, M. Coquery, N. Delorme, L. Garnero, L. Dherret, O. Geffard, and C. Lopes. One and multi-compartments toxico-kinetic modeling to understand metals ’

organotropism and fate in *Gammarus fossarum*. *Environment international*, 156(April):1–9, 2021. doi: 10.1016/j.envint.2021.106625.

[2] L. Petzold. Automatic selection of methods for solving stiff anf nonstiff systems of ODEs.pdf, 1983.

[3] L. Petzold and A. Hindmarsh. A systematized collection of ode solvers. *Report of*, 1997.

[4] K. Soetaert, T. Petzoldt, and R. W. Setzer. Solving differential equations in R: Package deSolve. *Journal of Statistical Software*, 33(9):1–25, 2010. doi: 10.18637/jss.v033.i09.

[5] J. Zhang, Q.-G. Tan, L. Huang, Z. Ye, X. Wang, T. Xiao, Y. Wu, W. Zhang, and B. Yan. Intestinal uptake and low transformation increase the bioaccumulation of inorganic arsenic in freshwater zebrafish. *Journal of Hazardous Materials*, 434(April):128904, 2022. ISSN 03043894. doi: 10.1016/j.jhazmat.2022.128904.
